# Multifunctional role of GPCR signaling in epithelial tube formation

**DOI:** 10.1101/2022.01.06.475238

**Authors:** Vishakha Vishwakarma, Thao Phuong Le, SeYeon Chung

## Abstract

Epithelial tube formation requires Rho1-dependent actomyosin contractility to generate the cellular forces that drive cell shape changes and rearrangement. Rho1 signaling is activated by G protein-coupled receptor (GPCR) signaling at the cell surface. During *Drosophila* embryonic salivary gland (SG) invagination, the GPCR ligand Folded gastrulation (Fog) activates Rho1 signaling to drive apical constriction. The SG receptor that transduces the Fog signal into Rho1-dependent myosin activation has not been identified. Here, we reveal that the Smog GPCR transduces Fog signal to regulate Rho kinase accumulation and myosin activation in the medioapical region of cells to control apical constriction during SG invagination. We also report on unexpected Fog-independent roles for Smog in maintaining epithelial integrity and organizing cortical actin. Our data supports a model wherein Smog regulates distinct myosin pools and actin cytoskeleton in a ligand-dependent manner during epithelial tube formation.

## INTRODUCTION

The formation of three-dimensional tubes by invagination from flat epithelial sheets is a fundamental process in forming organs, such as lungs and kidneys (Andrew and Ewald, 2010). The *Drosophila* embryonic salivary gland (SG) is a premier model system to study the mechanisms underlying epithelial tube morphogenesis (Chung et al., 2014; Girdler and Röper, 2014). The SG begins as a two-dimensional plate of cells on the embryo surface. Neither cell division nor cell death occurs once the SG cells are specified; all the morphogenetic changes arise by changes in cell shape and rearrangement.

A major cell shape change during SG invagination is apical constriction, wherein the apical side of epithelial cells shrinks while keeping the nearly constant volume (Lubarsky and Krasnow, 2003; Martin and Goldstein, 2014). During stage 11, SG cells begin to invaginate at the dorsal/posterior region of the placode through apical constriction (Fig. 1A). This leads to forming a narrow invagination pit through which all SG cells eventually internalize. Apical constriction is observed in both invertebrates and vertebrates, driving morphogenetic processes ranging from gastrulation in *Drosophila* and *Xenopus* to lens formation in the mouse eye to formation of tubular organs such as the *Drosophila* SG and chicken lungs (Kim et al., 2013; Myat and Andrew, 2000; Plageman et al., 2011; Sweeton et al., 1991). Defects in apical constriction result in defects in overall tissue shape, suggesting that coordinated apical constriction is required for final tissue architecture (Chung et al., 2017; Guglielmi et al., 2015; Izquierdo et al., 2018).

**Figure 1.**
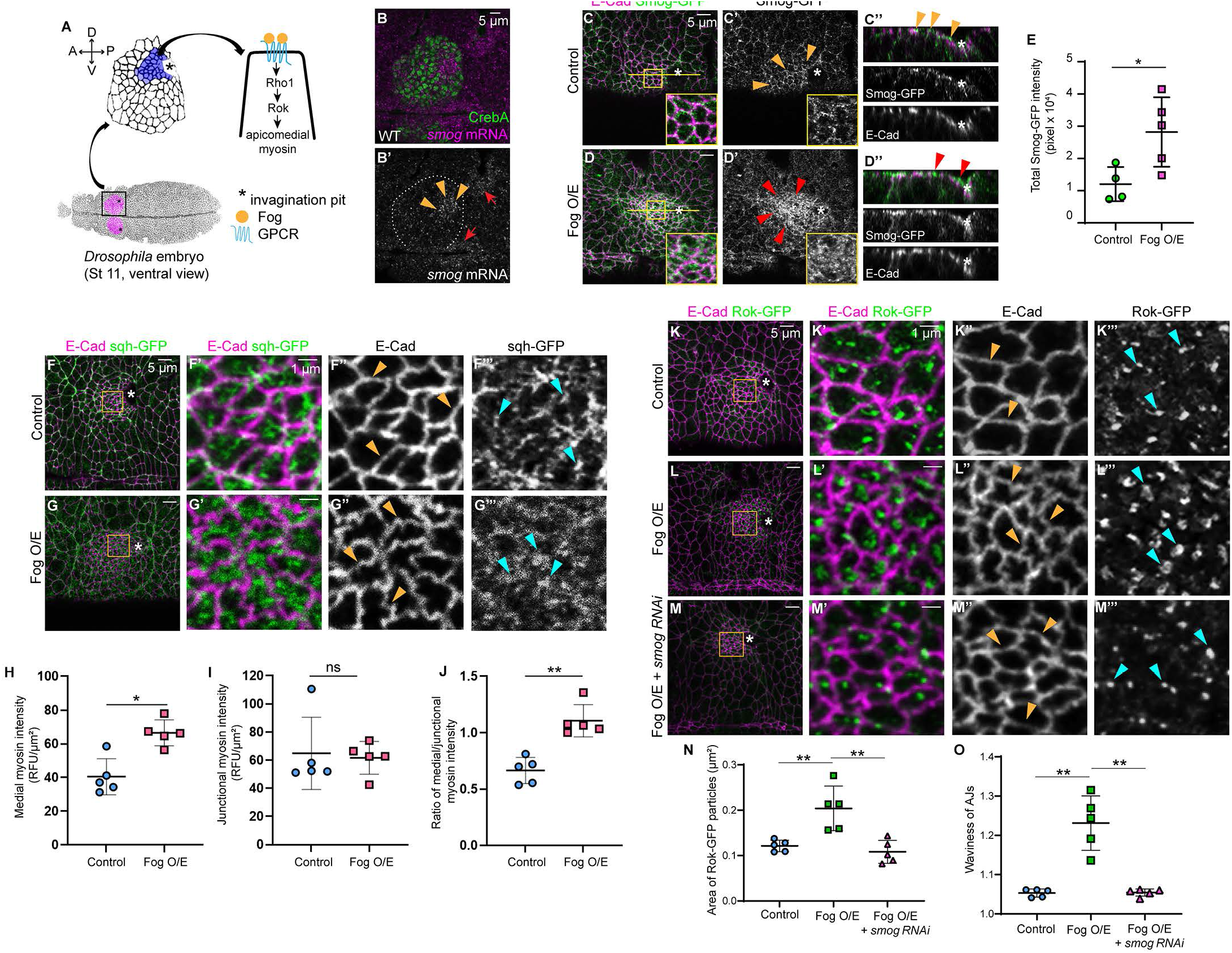
Smog transduces Fog signal to activate Rok and myosin in the SG. (A) Cartoon diagram showing SG placodes (magenta) in the ventral side of a stage 11 *Drosophila* embryo. Cells that undergo apical constriction near the invagination pit (asterisk) are shown in purple. In those cells, the Fog ligand (yellow) binds to its GPCR (blue) to activate medioapical myosin. In all stage 11 SG images, anterior is to the left, and dorsal is up. (B, B’) Fluorescent in situ hybridization of *smog* (magenta) in the WT SG. Arrowsheads, strong *smog* mRNA signals in apically constricting SG cells. Arrows, *smog* signals near the grooves and the ventral midline. Green, CrebA (SG nuclei). White lines, SG boundary. (C-D’’) Smog-GFP (green) signals in control (C-C’’) and Fog-overexpressing SGs (D-D’’). Magenta, E-Cad. Yellow arrowheads, puncta of Smog-GFP signals in the apical region near AJs in control SG cells. Rred arrowheads, considerably higher Smog-GFP signals in the entire apical domain of Fog-overexpressing SG cells. Insets, magnified images of yellow boxed regions. (C’’, D’’) z-sections across the yellow lines in C and D. (E) Quantification of total Smog-GFP intensity. n=4 SGs (control); 5 SGs (Fog overexpression). *p≤0.05 (Welch’s t-test). (F-G’’’) sqh-GFP (green) and E-Cad (magenta) signals in control (F-F’’’) and Fog-overexpressing (G-G’’’) SGs. (F’-G’’’) Magnified images of yellow boxed regions in F and G. Yellow arrowheads, AJs. Cyan arrowheads, sqh-GFP signals. (H-J) Quantification of medioapical myosin intensity (H), junctional myosin intensity (I), and the ratio of medioapical to junctional myosin intensity (J). ns, non-significant; *p≤0.05; **p≤0.01 (Welch’s t-test). (K-M’’’) Rok-GFP (green) and E-Cad (magenta) signals in SGs. K’-M’’’, magnified images of yellow boxed regions in K-M. Yellow arrowheads, AJs. Cyan arrowheads, Rok-GFP puncta. (N, O) Quantification of the area of Rok-GFP particles (N) and waviness of AJs (O). Asterisks, invagination pit. n=5 SGs, 75 cells for each genotype in H-J, N, and O. **p≤0.01 (Mann-Whitney U test).

Contractile actomyosin networks generate the cellular forces driving coordinated cell behaviors during epithelial morphogenesis (Gorfinkiel and Blanchard, 2011; Levayer and Lecuit, 2012; Munjal and Lecuit, 2014). Non-muscle myosin II (hereafter referred to as myosin) uses its motor activity to pull on actin filaments anchored to adherens junctions (AJs) to apply tension on the junction and neighboring cells. Three distinct pools of myosin generate the contractile forces required for SG invagination (Booth et al., 2014; Chung et al., 2017; Röper, 2012). Medioapical myosin forms in the center of the apical side of the SG cell and drives apical constriction (Booth et al., 2014; Chung et al., 2017; Röper, 2012). Junctional myosin, associated with AJs of SG cells, is involved in cell intercalation (Sanchez-Corrales et al., 2018). A supracellular myosin cable surrounding the entire SG placode generates the tissue-level compressing force (Chung et al., 2017; Röper, 2012). From studies in the past several years by us and others, we are beginning to understand how each myosin pool is created and regulated during SG invagination (Booth et al., 2014; Chung et al., 2017; Le and Chung, 2021; Röper, 2012; Sanchez-Corrales et al., 2018; Sidor et al., 2020).

Localized activation of the RhoA (Rho1 in *Drosophila*) GTPase is key to creating and/or polarizing contractile myosin structures in epithelial cells (Blanchard et al., 2010; Dawes-Hoang et al., 2005; Martin et al., 2009; Mason et al., 2013). Studies in *Drosophila* have identified the Folded gastrulation (Fog) pathway as a key signaling pathway regulating Rho1 signaling during epithelial morphogenesis (Manning and Rogers, 2014). During *Drosophila* gastrulation, two G protein-coupled receptors (GPCRs), Smog (ubiquitously expressed) and Mist (mesoderm-specific), respond to Fog to activate heterotrimeric G proteins, Rho1, and Rho kinase (Rok) to regulate myosin contractility (Costa et al., 1994; Dawes-Hoang et al., 2005; Kerridge et al., 2016; Kölsch et al., 2007; Manning et al., 2013; Mason et al., 2013; Parks and Wieschaus, 1991). In our previous study in the SG, we also revealed a tissue-specific regulation of Fog signaling to activate myosin contractility: the Fork head (Fkh) transcription factor regulates SG upregulation of *fog*, which promotes accumulation of Rok and myosin specifically in the medioapical region of SG cells to drive clustered apical constriction (Chung et al., 2017; Fig. 1A). Yet, it is unknown how the Fog signal is sensed and transduced into downstream Rho1 signaling during epithelial tube formation.

Studies in early *Drosophila* embryos have suggested a role of GPCR signaling as a common module for Rho1 activation in different subcellular regions of the cell. In the mesoderm, Smog and Mist function together to activate myosin in the medioapical region of cells to drive apical constriction (Kerridge et al., 2016). In the ectoderm, where no apical constriction occurs, Smog is required for junctional myosin activation to drive cell intercalation (Kerridge et al., 2016). Recent work further has shown that distinct RhoGEFs activate myosin contractility in the apical and junctional domain of epithelial cells under the control of specific G proteins in early *Drosophila* embryos (Garcia De Las Bayonas et al., 2019). However, whether all three myosin pools in the SG are regulated by independent mechanisms or by a common module during epithelial tube formation remains unclear.

The contractile force of the actomyosin networks causes an increase of hydrostatic pressure in the cytoplasm (Charras et al., 2005). During epithelial morphogenesis, cells maintain the surface integrity by proper organization of actin at the cell cortex. Loss of organized cortical actin networks leads to the formation of blebs, spherical protrusions of the plasma membrane (Charras, 2008). During *Drosophila* gastrulation, blebs occasionally arise during apical constriction of mesodermal cells; these blebs correlate with apical F-actin holes (Costa et al., 1994; Jodoin et al., 2015). Disruption of cortical actin by latrunculin B, which sequesters globular actin (G-actin) and prevents filamentous actin (F-actin) assembly, enhances blebbing in epithelial cells in early *Drosophila* embryos (Kanesaki et al., 2013). Mutations in heterotrimeric G proteins also enhance blebbing (Kanesaki et al., 2013), suggesting a role of G proteins in stabilizing cortical actin. The mechanisms of cortical actin regulation during epithelial tube formation are not well understood.

Here, we reveal multiple roles of Smog GPCR in regulating actomyosin networks during *Drosophila* SG tube formation. We show that the GPCR Smog transduces Fog signal to regulate myosin contractility in the medioapical region of SG cells to drive apical constriction during SG invagination. Our study supports a model wherein Smog regulates medioapical and junctional/supracellular myosin pools in a Fog-dependent and -independent manner, respectively. We further reveal new roles of Smog in regulating the microtubule networks, key apical and junction proteins, and cortical actin organization during SG invagination, suggesting multifunctional roles of GPCR signaling during epithelial tube formation.

## RESULTS

### Smog transduces Fog signal in the SG

To test the role of Smog in transducing Fog signal during SG invagination, we first examined *smog* transcripts and Smog protein levels in the developing SG. Consistent with the ubiquitous expression of *smog* (Kerridge et al., 2016), fluorescent in situ hybridization revealed low levels of *smog* mRNA expression in the entire embryo, including the SG. Strong *smog* mRNA signals were observed in SG cells near the invagination pit (Fig. 1B, B’), potentially due to the shrinking apical domain in those cells. We also examined Smog protein localization in the SG using a functional Smog-GFP fusion protein expressed under the control of the *sqh* promoter (Kerridge et al., 2016). Smog-GFP signals were detected as small punctate structures enriched near AJs in SG cells (Fig. 1C-C’’; AJs were marked with the key AJ protein E-Cadherin (E-Cad). Importantly, Smog-GFP signals were dramatically increased in the entire apical domain of Fog-overexpressing SGs (Fig. 1D-D’’; quantification in Fig. 1E), suggesting that Smog is recruited to the apical domain of SG cells by overproduced Fog.

To test whether Smog transduces Fog signal in the SG to regulate myosin contractility, we performed a genetic suppression test. We hypothesized that if Smog is a SG receptor for Fog, knocking down *smog* in the SG should suppress the gain-of-function effect of Fog. We used sqh-GFP, a GFP-tagged version of myosin regulatory light chain (Royou et al., 2004), and Rok-GFP, a GFP-tagged Rok fusion protein (Abreu-Blanco et al., 2014), as readouts of Fog signaling. sqh-GFP signals were significantly increased in the medioapical region of Fog-overexpressing SG cells (compare Fig. 1G’, G’’’ to Fig. 1F’, F’’’; quantification in Fig. 1H) but not at AJs (Fig. 1I), suggesting increased medioapical myosin by Fog overexpression. Fog-overexpressing SG cells showed highly distorted AJs (Fig. 1G’’) compared to control SG cells (Fig 1F’’), suggesting increased pulling forces due to increased medioapical myosin in Fog-overexpressing SG cells. Consistent with our finding that Fog signal promotes accumulation of Rok in the medioapical region of SG cells (Chung et al., 2017), quantification of areas occupied by Rok-GFP puncta revealed overaccumulated Rok-GFP signals in the medioapical region of Fog-overexpressing SG cells (compare Fig. 1L’, L’’’ to Fig. 1K’, K’’’; quantification in Fig. 1N). Knockdown of *smog* in the SG using RNA interference (RNAi) and the SG-specific *fkh-Gal4* driver (Henderson and Andrew, 2000) both suppressed the medioapical accumulation of Rok-GFP signals and reduced the magnitude of AJ distortion (Fig. 1M-M’’’; quantification in 1N, O). These data suggest that Smog transduces Fog signal in the SG to facilitate Rok and myosin accumulation in the medioapical region of SG cells.

### Smog-transfected *Drosophila* S2 cells contract upon Fog signal

To further test the role of Smog in transducing Fog signal to regulate myosin contractility, we performed an in vitro cell contraction assay using *Drosophila* S2 cells. S2 cells do not express Fog receptors, and therefore, do not respond to Fog in the normal culture condition (Manning et al., 2013; Fig. 2A-B’). As shown in a previous study (Manning et al., 2013), S2 cells transfected with Mist, the mesoderm-specific receptor for Fog, contracted robustly when cultured in Fog-containing media (Fig. 2C-D’). Since Smog was not detectable in *Drosophila* S2 cells (modENCODE Cell Line Expression Data; Flybase), we tested if S2 cells transfected with Smog also contract upon Fog treatment. Indeed, Smog-transfected cells showed a significantly higher percentage of contraction upon Fog treatment compared to non-transfected S2 cells (Fig. 2A-B’, E-F’; quantification in Fig. 2G). Thus, Smog can transduce Fog signal to regulate cellular contractility both in the SG and in vitro.

**Figure 2.**
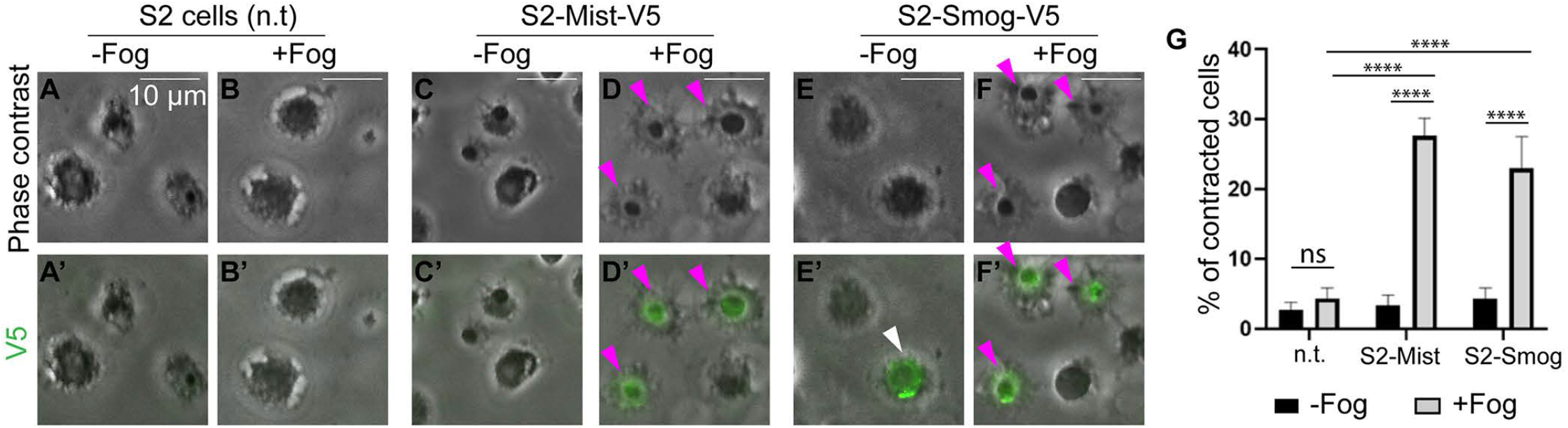
Smog-transfected S2 cells contract in response to Fog. (A-F’) S2 cells cultured in the absence (A, C, E) or presence (B, D, F) of Fog. (A-B’) Non-transfected. (C-D’) Mist-transfected. (E-F’) Smog-transfected. A-F, phase contrast; A’-F’, merged images of phase contrast and V5 signals for transfected cells (green). White arrowhead in E’, a Smog-transfected S2 cell not contracting in the absence of Fog. Magenta arrowheads, contracting cells. (G) Quantification of percentage of contracting cells among the transfected cells. n=∼100 cells per condition from three independent experiments. ****p≤0.0001 (Two-way ANOVA with Tukey’s multiple comparison post hoc analysis).

### Smog is required for apical constriction in the SG and epithelial morphogenesis during embryogenesis

To test the role of Smog in apical constriction of SG cells as a Fog receptor, we knocked down *smog* in the SG (using *fkh-Gal4*) using RNAi and compared apical areas of SG cells to those in *fog* mutants. Two independent short hairpin RNAi lines, *TRiP*.*HMC03192* (a stronger line; Fig. 3B, B’; also used for genetic suppression assay in Fig. 1M-M’’’) and *TRiP*.*GL01473* (a weaker line; Fig. S1G, G’), were used. *smog* knockdown using either line reduced Smog-GFP signals in the SG, with a stronger effect with *TRiP*.*HMC03192*, confirming the effect of the RNAi lines (Fig. S1A-D). Cell segmentation analysis revealed that SG cells with smaller apical areas showed a less coordinated spatial distribution in *smog* RNAi SGs compared to control (Fig. 3A-B’; Fig. S1F-G’), similar to the previously reported *fog* mutant phenotype (Chung et al., 2017; Fig. S1E, E’). However, quantification of percentage and cumulative percentage of cells of different apical areas revealed different behaviors between *fog* and *smog*. In *fog* mutants, the number of constricting cells is not significantly different from wild type (WT), but cells with small apical areas are dispersed across the entire SG placode (Chung et al., 2017). On the other hand, SGs with *smog* knockdown with either RNA line showed larger apical areas compared to control (Fig. 3G; Fig. S1H). The difference in apical constriction defects between *fog* mutants and *smog* knockdown suggests a potential Fog-independent role of Smog.

**Figure 3.**
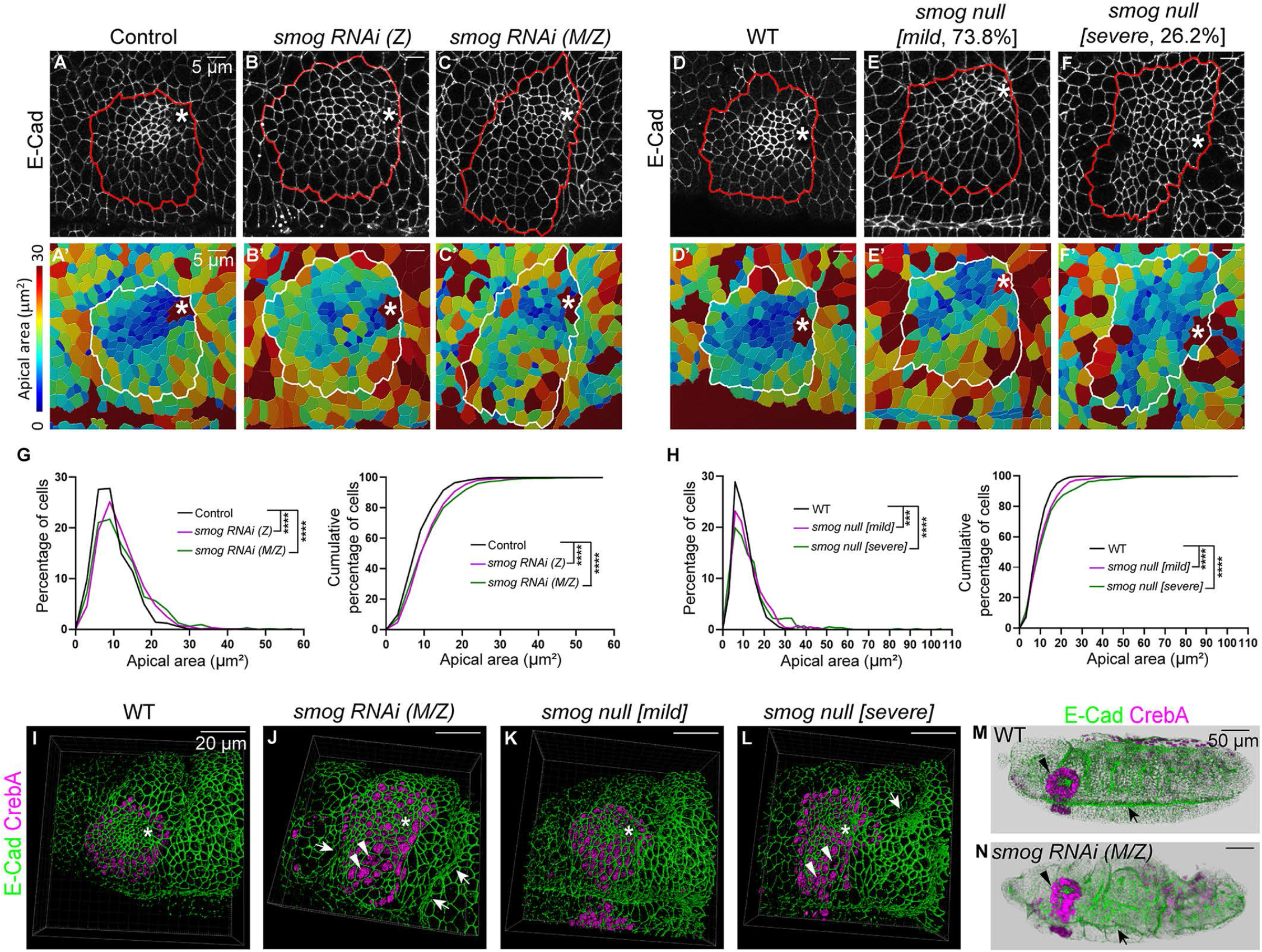
*smog* knockdown and loss result in apical constriction defects in the SG and disrupted epithelial morphology. (A-F’) Confocal images of stage 11 SGs stained for E-Cad (A-F) and corresponding heat maps of apical areas (A’-F’). *TRiP*.*HMC03192* and *TRiP*.*GL01473* are used for zygotic and M/Z knockdown of *smog*, respectively. Red and white lines, SG boundary. (G, H) Quantification of percentage (left) and cumulative percentage (right) of cells with different apical areas. Mann-Whitney U test (for percentage of cells) and Kolmogorov-Smirnov test (for cumulative percentage of cells). n=8 SGs (control, 902 cells; *smog* RNAi (Z), 823 cells; *smog* RNAi (M/Z), 881 cells; WT, 892 cells; *smog* null [mild], 859 cells); 6 SGs (*smog* null [severe], 672 cells). ***p<0.001; ****p<0.0001. (I-L) 3D reconstruction of confocal z-sections of SGs (Imaris) for WT (I), M/Z knockdown of *smog* (J), *smog* null [mild] (K), and *smog* null [severe] (L). Green, E-Cad. Magenta, CrebA. Asterisks, invagination pit. Arrows in J and L, exaggerated grooves. Arrowheads in J and L, enlarged SG cells. (M, N) Tile scan confocal images of stage 11 embryos in WT (M) and M/Z knockdown of *smog* (N). Green, E-Cad. Magenta, CrebA. Arrows, the ventral midline. Arrowheads, SGs.

Since *smog* has a strong maternal contribution (Flybase Developmental RNA-Seq), we also knocked down *smog* both maternally (using *matα-Gal4*; Häcker and Perrimon, 1998) and zygotically in the SG (using *fkh-Gal4*) and observed slightly more severe apical constriction defects in SG cells (hereafter referred to as *smog* M/Z knockdown; Fig. 3C, C’; quantification in 3G). Maternal knockdown of *smog* using the stronger RNAi line (*TRiP*.*HMC03192*) resulted in severe defects in egg-laying. Therefore, all M/Z knockdown experiments were performed using the weaker line (*TRiP*.*GL01473*). Consistent with the ubiquitous expression of *smog* and its role in epithelial morphogenesis in the early *Drosophila* embryo (Kerridge et al., 2016), *smog* M/Z knockdown resulted in a slight disruption of the overall embryonic morphology, including defects in the head region, a wavy embryo surface, and an irregular ventral midline (Fig. 3M, N). Unlike the circular SG placode of WT, *smog* M/Z knockdown also resulted in a SG placode elongated along the dorsal/ventral axis (compare Fig. 3A, A’ to 3C, C’). 3D reconstruction of confocal images revealed disrupted epithelial morphology in *smog* M/Z knockdown, with wide and wavy grooves, enlarged epithelial cells, and an elongated SG placode (Fig. 3I, J).

*smog* zygotic null mutants (Kerridge et al., 2016) had a range of defects similar to those observed with *smog* knockdown. 73.8% of embryos (59/80) showed relatively normal embryonic morphology with mild apical constriction defects in SG cells, like *smog* zygotic knockdown (classified as ‘*smog* null [mild]’; Fig. 3D-E’, K; quantification in 3H); 26.2% of embryos (21/80) showed a disrupted embryonic morphology with SGs elongated along the dorsal/ventral axis (Fig. 3F, F’, L) and more severe apical constriction defects (Fig. 3H), like *smog* M/Z knockdown (classified as ‘*smog* null [severe]’). All embryos at late stages (stages 15-17) *smog* null mutants showed relatively normal embryonic morphology (Fig. S2A, B), suggesting that severely defective embryos with *smog* loss do not survive/develop past stage 14. Notably, these embryos have relatively normal internalized SGs (Fig. S2E-G), except for rare cases of crooked SG morphology (Fig. S2C, D). Altogether, these data suggest that Smog is required for coordinated apical constriction during SG invagination and proper epithelial morphogenesis during embryogenesis.

### Smog regulates medioapical and junctional myosin in a Fog-dependent and -independent manner in the SG

We next examined myosin levels and distribution in the SGs in *smog* knockdown or *smog* mutants using sqh-GFP. Medioapical myosin is the only myosin pool defective in the SGs of *fog* mutants (Chung et al., 2017; Fig. S3A-E). Consistent with the idea that Smog transduces Fog signal in the SG, *smog* knockdown (either zygotic or M/Z knockdown) and *smog* loss resulted in a significant reduction in the intensity of medioapical myosin in SG cells near the invagination pit, compared to control (Fig. 4A-D’’; quantification in 4I). Moreover, compared to clear web-like structures of medioapical myosin in control SG cells (Fig. 4A-A’’), SG cells in *smog* knockdown or *smog* null mutants showed reduced areas of sqh-GFP puncta (Fig. 4L), with dispersed myosin along the entire apical surface in *smog* M/Z knockdown SG cells (Fig. 4C-C’’).

**Figure 4.**
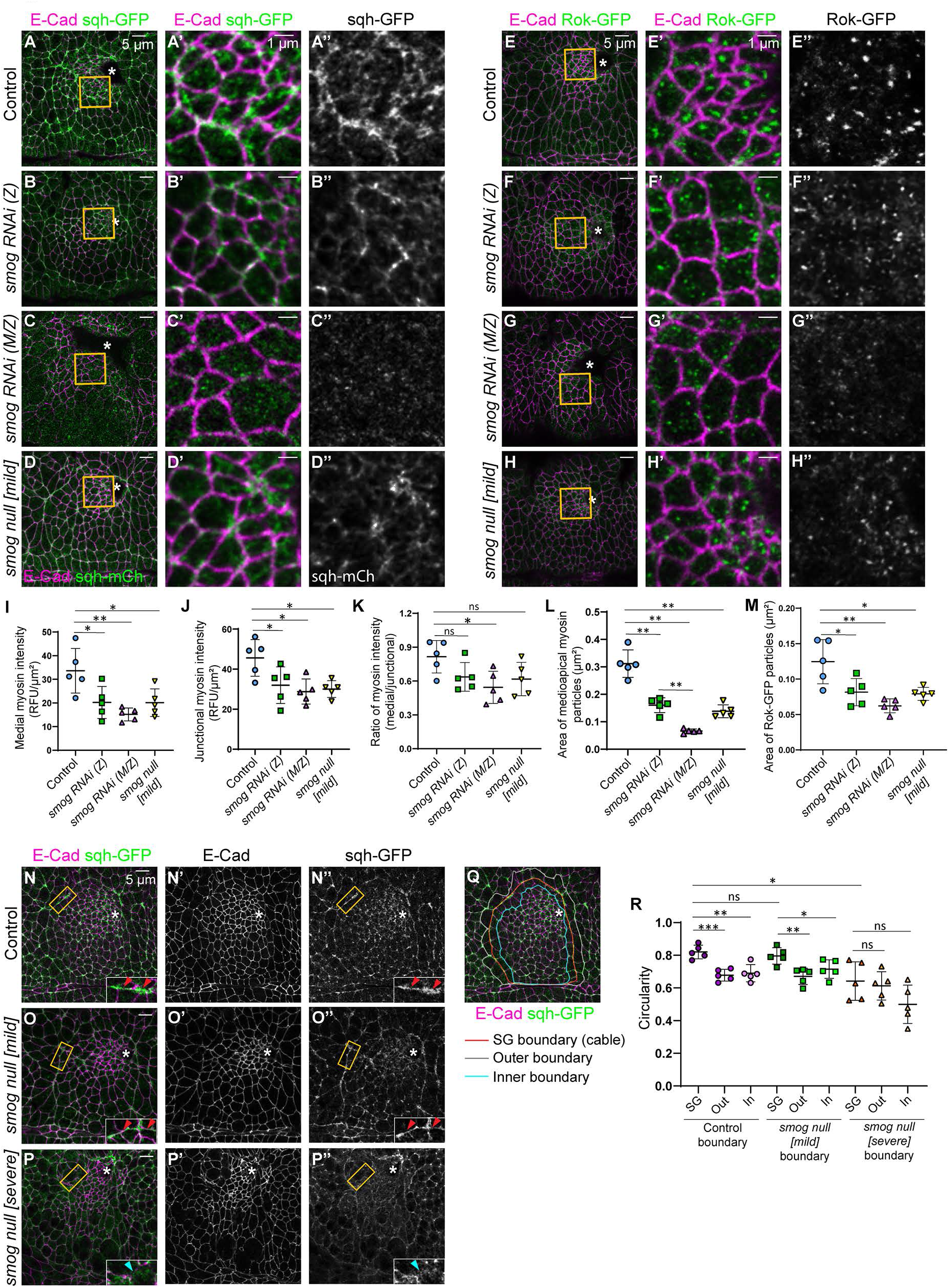
*smog* regulates all three pools of myosin in the SG in a Fog-dependent and -independent manner. (A-D’’) Confocal images of stage 11 SGs with sqh-GFP (green) and E-Cad (magenta) signals. (A’-D’’) Higher magnification of the yellow boxed region in A-D. (E-H’’) Confocal images of SGs with Rok-GFP (green) and E-Cad (magenta) signals. (E’-H”) Magnified images of the yellow boxed region in E-H. (I-M) Quantification of medioapical myosin intensity (I), junctional myosin intensity (J), ratio of medial and junctional myosin intensity (K), area of medioapical myosin particles (L), and area of Rok-GFP particles (M). ns, non-significant; *p≤0.05; **p≤0.01 (Welch’s t-test for I-K; Mann-Whitney U test for L and M). n=5 SGs, 100 cells (I-L) and 5 SGs, 75 cells (M) for each genotype. (N-P’’) Confocal images of SGs with sqh-GFP (green) and E-Cad (magenta) signals. Insets, magnified views of the SG boundary. Red arrowheads in N, N’’, O and O’’, high myosin intensity at the SG boudary in control and *smog* null [mild] embryos. Cyan arrowheads in P and P’’, low myosin signals at the SG boundary in the *smog* null [severe] embryo. (Q) Circularity of the SG placode boundary. An example of a WT SG stained for E-Cad (magenta) and sqh-GFP (green) with the SG boundary (red line) and the boundaries one cell row outside (gray line) and inside (blue line) the SG boundary. (R) Quantification of circularity of the SG, outer, and inner boundaries. Asterisks, invagination pit. n=5 SGs for each genotype. ns, non-significant; *p≤0.05; **p≤0.01; ***p≤0.001 (Welch’s t-test).

Interestingly, the intensity of junctional myosin was also significantly reduced in SG cells with *smog* knockdown or *smog* loss, compared to control (Fig. 4A-D’’, quantification in 4J). This phenotype is distinct from what was observed in *fog* mutants, where junctional myosin was unaffected in SG cells (Chung et al., 2017; Fig. S3D). The ratio of medioapical to junctional myosin intensity was significantly reduced in *smog* M/Z knockdown but not in *smog* zygotic knockdown or *smog* null [mild] SGs (Fig. 4K). We conclude that Smog is required for maintaining normal levels of both medioapical and junctional myosin in the SG in a Fog-dependent and -independent manner, respectively.

We also tested Rok distribution in SG cells in *smog* RNAi and *smog* null mutants using Rok-GFP. Whereas Rok-GFP formed large punctate structures in the medioapical region in control SG cells (Fig. 4E-E’’), Rok-GFP failed to accumulate and was dispersed along the entire apical domain in SG cells in *smog* knockdown and *smog* null mutants (Fig. 4F-H’’; quantification in 4M), similar to the dispersed Rok-GFP distribution observed in *fog* mutant SG cells (Chung et al., 2017). These data suggest that Smog is required for Rok accumulation in the medioapical domain of SG cells.

### *smog* loss leads to defects in the tissue-level myosin cable

Reduced myosin levels and the elongated SG placode in *smog* null [severe] embryos led us to test the third pool of myosin in the SG, the supracellular myosin cable surrounding the SG placode. Consistent with the idea that this myosin cable is under tension (Chung et al., 2017; Röper, 2012), WT SGs showed a relatively smooth tissue boundary (Fig. 4N-N’’). *smog* null [mild] embryos also maintained a smooth SG boundary comparable to WT (Fig. 4O-O’’). However, SGs in *smog* null [severe] embryos showed an irregular boundary and a less clear myosin cable (Fig. 4P-P’’).

We calculated the circularity of the SG placode as a measure of smoothness and tension of the SG boundary. The circularity of the WT placode was significantly higher than the circularity of one cell row inside (inner boundary) or outside of the SG placode (outer boundary), suggesting a higher tension of the myosin cable in WT (Fig. 4Q, R). The circularity of the placode of *smog* null [mild] embryos was comparable to WT (Fig. 4R). However, the circularity of the SG placode in *smog* null [severe] embryos was significantly lower than that of WT and was not statistically significant from the inner or outer boundaries, suggesting a lack of tension at SG boundary in *smog* null [severe] mutants (Fig. 4R). Consistent with our previous study (Chung et al., 2017), *fog* mutants did not show defects in the myosin cable (Fig. S3F-H), even in the severely twisted embryos with extra grooves and folds. Overall, our data suggest that, unlike *fog*, the loss of *smog* affects all myosin pools in the SG, including the myosin at the SG boundary, leading to reduced tension at the SG boundary and defects in generating the compressing force.

### Smog is required for proper localization of apical and junctional components and microtubule networks in epithelial cells

Since we observed a range of defects in whole embryo morphology in *smog* mutants, we explored the potential cellular basis. In stage 11 *smog* null [severe] embryos, we occasionally observed large areas in the epidermis where E-Cad (Fig. 5B, B’; compare to WT in Fig. 5A, A’) and the apical determinant protein Crumbs (Crb) (Fig. 5D, D’; compare to WT in Fig. 5C, C’) signals were absent or significantly reduced. Relatively strong signals of the cell membrane marker Gap43-mCherry (Martin et al., 2010) were observed in these regions (Fig. 5E-E’’), suggesting that the cell membrane is still intact. Cells in these regions had enlarged apical areas compared to neighboring cells with strong E-Cad signals (Fig. 5E-E’’), suggesting reduced epithelial tension and/or integrity in cells with reduced E-Cad levels. Similarly, in late-stage (stage 14) embryos, Crb levels were significantly reduced in large areas; in the most severe cases, Crb was dispersed in the entire epidermis (Fig. S4B, B’). These data suggest that Smog is required for maintaining epithelial polarity and junctional integrity by regulating E-Cad and Crb levels or distribution.

**Figure 5.**
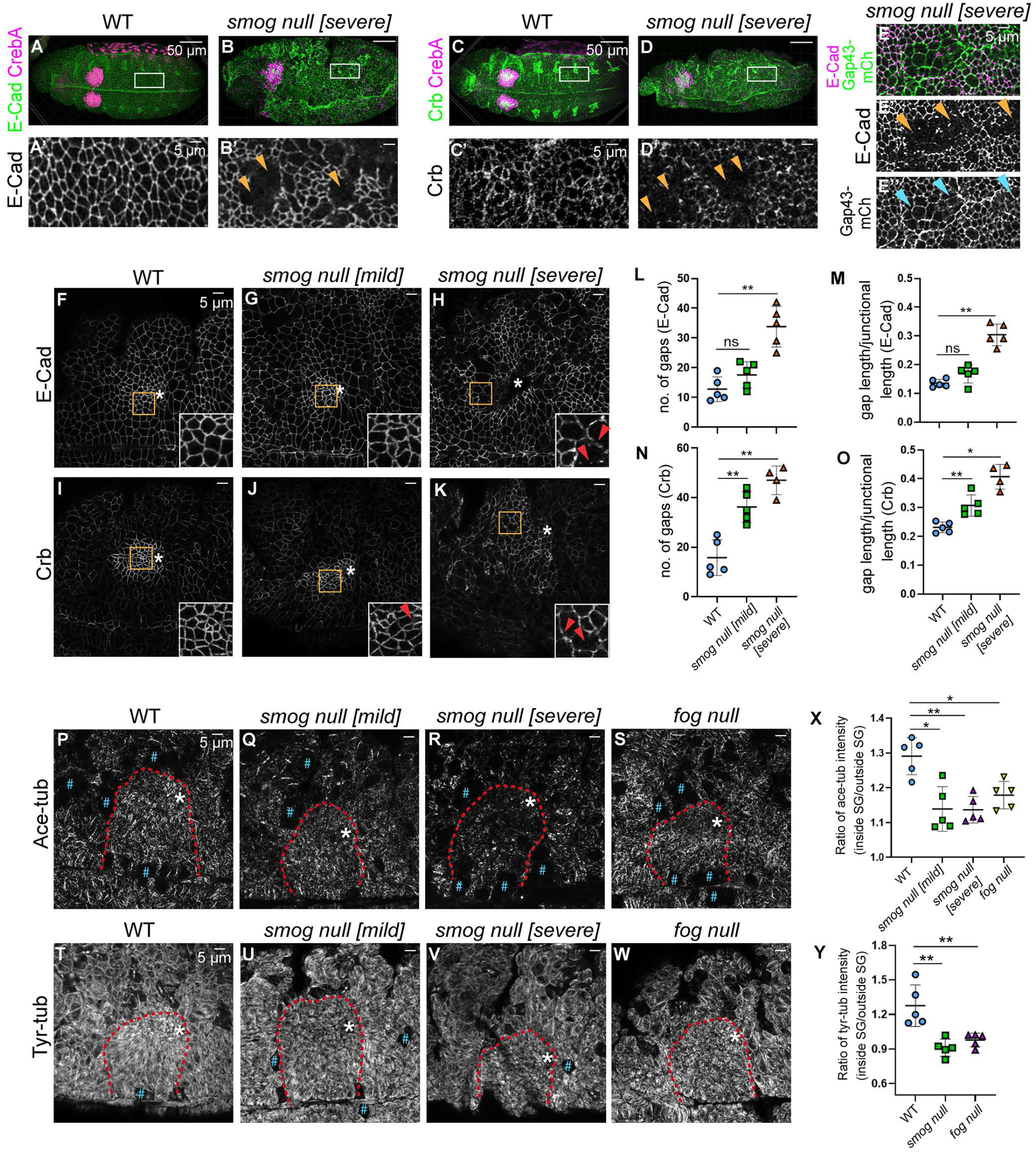
Smog is required for maintaining epithelial integrity and the microtubule networks. (A-D’) Confocal images of stage 11 embryos stained for E-Cad (green in A, B), Crb (green in C, D) and CrebA (magenta). (A’-D’) Magnified view of the white boxed region in A-D. Arrowheads in B’ and D’, E-Cad (B’) and Crb (D’) signals are absent in regions of the epidermis in *smog* mutants. (E) E-Cad (magenta) and Gap43-mCh (green) signals in the epidermis in the *smog* null [severe] embryo. The cell membrane is intact (cyan arrowheads in E’’) in regions where E-Cad signals are absent (yellow arrowheads in E’). (F-K) E-Cad (F-H) and Crb (I-K) signals in SGs in WT (F, I), *smog* null [mild] (G, J), and *smog* null [severe] (H, K) embryos. Insets, magnified view of yellow boxed region in the SG. Arrowheads, gaps in E-Cad or Crb signals. (L-O) Quantification of the number of gaps in E-Cad (L) and Crb (N) signals and the ratio of gap length to total junctional length for E-Cad (M) and Crb (O). n=5 SGs, 50 cells per genotype. ns, non-significant; *p≤0.05; **p≤0.01 (Mann-Whitney U test). (P-W) Ace-tub (P-S) and Tyr-tub (T-W) signals in the SG and the surrounding epithelia. Red dotted lines, SG boundary. (X, Y) Quantification of the ratio of Ace-tub (X) and Tyr-tub (Y) intensity in SG cells to the intensity in cells outside the SG placode. n=5 SGs for each genotype. *p≤0.05; **p≤0.01 (Mann-Whitney U test). Asterisks, invagination pit; Pound sign, dividing cells.

To better understand the basis of the reduced E-Cad and Crb in *smog* mutants, we examined *smog* null mutant SG cells at higher magnification. Even in SGs that do not contain large patches of E-Cad/Crb loss, E-Cad/Crb signals were discontinuous, with small gaps in signal along the cell boundary (Fig. 5F-O). Such gaps were observed in both *smog* null [mild] and *smog* null [severe] embryos. Although the number and the length of gaps in E-Cad signals in *smog* null [mild] were comparable to those of WT, they were significantly higher in *smog* null [severe] compared to WT (Fig. 5G, H; quantification in 5L, M). The number and the length of gaps in Crb signals in SG cells were significantly higher in both mild and severe *smog* null mutant embryos compared to WT (Fig. 5J, K; quantification in 5N, O). These data suggest a role of Smog in regulating Crb and E-Cad levels and continuity during epithelial morphogenesis, including the SG.

Anisotropic localization of Crb at the SG boundary has been suggested to drive the formation of supracellular myosin cable in SG (Röper, 2012). Consistent with this, in control SGs, Crb levels were higher in inside junctions that do not contribute to the myosin cable, whereas myosin was highly enriched at the junction forming the cable at the SG boundary (Fig. S4C-E’’’). However, in *smog* null [severe] SGs, the overall Crb intensity was quite low, and Crb did not show an anisotropic localization at the SG boundary (Fig. S4F-H’’’). This data suggests that *smog* loss results in the reduction of both myosin and Crb, failing to form the supracellular myosin cable at the SG boundary.

Microtubules are required for forming and maintaining apical myosin structures (Booth et al., 2014; Ko et al., 2019) and for transporting several key apical and junctional proteins in *Drosophila* tubular organs, including E-Cad and Crb (Le and Chung, 2021; le Droguen et al., 2015). We recently showed that microtubule-dependent intracellular trafficking has an important role in regulating apical constriction and medioapical myosin during SG invagination (Le and Chung, 2021). To examine the microtubule networks in the SG of WT and *smog* mutant embryos, we stained with antibodies against acetylated α-tubulin (Ace-tub), a marker of stable, long-lived microtubules, and tyrosinated α-tubulin (Tyr-tub), a marker of dynamic, short-lived microtubules (Westermann and Weber, 2003). Compared to WT, both Ace-tub and Tyr-tub signals were reduced in *smog* mutants (compare Fig. 5Q, R, U, V to Fig. 5P, T), with a dramatic reduction in Ace-tub levels in *smog* null [severe] embryos (Fig. 5R), suggesting disrupted microtubule networks in *smog* mutants. The reduced Ace-tub and Tyr-tub signals were more prominent in the SG compared to the surrounding epithelial cells; the ratio of Ace-tub and Tyr-tub intensity in SG cells to the intensity in cells outside the SG was significantly decreased in *smog* mutants compared to WT (Fig. 5X, Y). A slight but less prominent reduction in Ace-tub and Tyr-tub signals was also observed in the SG in *fog* mutants (Fig. 5S, W; quantification in 5X, Y). Overall, this finding suggests that loss of *smog* could affect apical pools of myosin, Crb and E-Cad through effects on microtubule abundance, distribution and/or polarity.

### Smog is required for cortical actin organization in the SG

We observed numerous blebs in the apical membrane of SG cells in *smog* null [severe] mutants, which were most prominent near the invagination pit (compare Fig. 6A-A’’’ to Fig. 6B-B’’’). Such blebs were not observed in SGs in WT, *smog* null [mild], or *fog* mutant embryos (Fig. S5A-C’’). Rok-GFP was enriched in many of these blebs (Fig. 6B-B’’’), consistent with Rok’s recruitment to retracting blebs (Aoki et al., 2016). In *smog* null [severe] embryos, blebs were also occasionally observed in cells outside the SG placode (Fig. S5D-E’’). During *Drosophila* gastrulation, bleb formation correlates with cortical F-actin holes (Jodoin et al., 2015), and disruption of cortical actin enhances blebbing (Kanesaki et al., 2013). We therefore tested if Smog affects the cortical actin network during apical constriction in the SG. Indeed, phalloidin staining revealed that whereas strong F-actin signals were observed in the apical domain of SG cells near the invagination pit in WT, F-actin was reduced in SGs in *smog* mutants (Fig. 6C-E).

**Figure 6.**
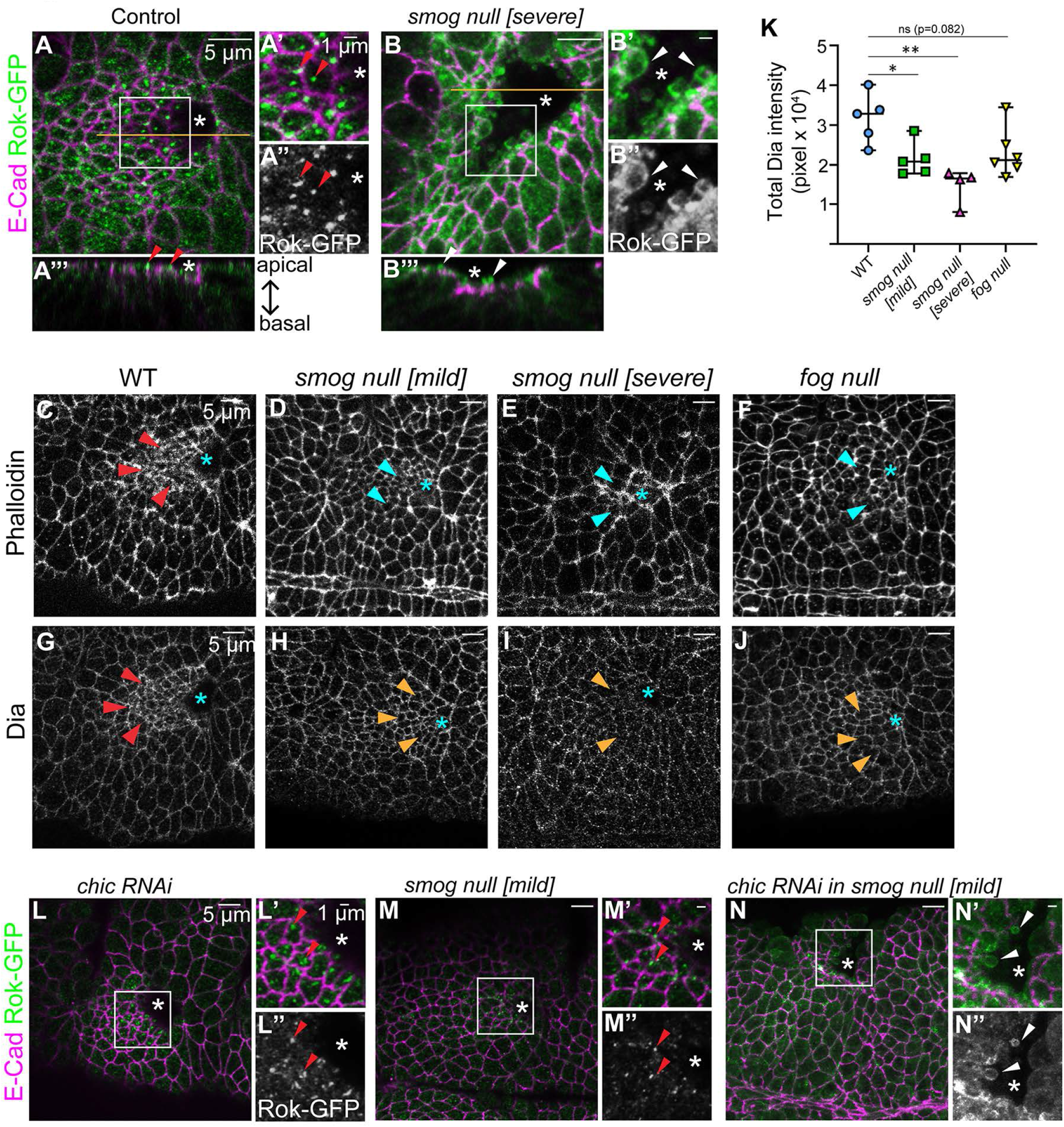
Smog is required for maintaining cortical actin networks during SG invagination. (A-B’’’) Rok-GFP (green) and E-Cad (magenta) in stage 11 SGs in control (A-A’’’) and *smog* null [severe] (B-B’’’) embryos. (A’-B’’) Higher magnification of the white boxed regions in A and B. (A’’’, B’’’) x-z sections across the yellow lines in A and B. Red arrowheads, normal Rok-GFP signals accumulated in the medioapical region of control SG cells. White arrowheads, blebs in SG cells in *smog* mutants enriched with Rok-GFP signals. (C-J) SGs stained for phalloidin (C-F) and Dia (G-J). Compared to strong phalloidin and Dia signals in WT (red arrowheads in C, G), phalloidin and Dia levels are reduced in *smog* and *fog* mutants (cyan arrowheads in D-F, yellow arrowheads in H-J). (K) Quantification of total Dia intensity in the SG placode. n=5 SGs for control and *smog* null [mild], n=4 SGs for *smog* null [severe], n=6 SGs for *fog* null. *p≤0.05; **p≤0.01 (Mann-Whitney U test). (L-N’’) Rok-GFP (green) and E-Cad (magenta) in *chic* RNAi in the wild-type (L-L’’), *smog* null [mild] (M-M’’), and *chic* RNAi in *smog* mutant background (N-N’’). (L’-N’’) Higher magnification of the white boxed regions in L-N. Red arrowheads in L’, L’’, M’, and M’’, normal Rok-GFP signals in *chic* RNAi and smog null [mild] SGs. White arrowheads in N’ and N’’, membrane blebs with enriched Rok-GFP signals in *smog* null [mild] SG cells with *chic* RNAi. Asterisks, invagination pit.

Apical F-actin formation in *Drosophila* tubular organs requires the formin family actin nucleator Diaphanous (Dia) (Massarwa et al., 2009; Rousso et al., 2013). Since the apical localization and activity of Dia are critical for restricting F-actin formation to the correct membrane domain, we tested the localization and levels of Dia in the SG in *smog* mutants. Similar to reduced F-actin (Fig. 6C-E), we observed reduced Dia signals in the apical domain of SG cells in *smog* mutants, compared to WT (Fig. 6G-I). Quantification of Dia levels revealed that the overall Dia intensity is reduced in the SG placode in *smog* mutants, with a stronger effect in the *smog* null [severe] than *smog* null [mild] embryos (Fig. 6K). SGs in *fog* mutants appeared to show a slight reduction of F-actin (Fig. 6F), but Dia levels in *fog* mutants were comparable to WT (Fig. 6J, K). Our data suggest that Smog is required for the cortical actin networks during SG invagination, at least in part via modulating Dia levels.

We next tested if the reduction in actin levels in *smog* mutants is due to defects in actin turnover. To test this, we modulated the level of actin regulators in the SG in *smog* mutants and assayed the blebbing phenotype. Using RNAi, we knocked down *chickadee* (*chic*), which encodes *Drosophila* profilin, a protein that increases F-actin by promoting actin polymerization (Cooley et al., 1992), in the *smog* mutant background. If there is a shift from F-actin to monomeric G-actin in *smog* mutants, we expect enhancement of the disorganized actin phenotype and bleb formation with a reduced *chic* level. Blebs were not observed in *smog* null [mild] embryos or in SGs knocked down for *chic* in the otherwise wild-type background (Fig. 6L-M’’). However, when *chic* was knocked down in the SG in the *smog* mutant background, we observed enhanced blebbing even in SG cells in *smog* null [mild] embryos (Fig. 6N-N’’). Since profilin promotes formin-mediated actin filament assembly (Romero et al., 2004; Zweifel and Courtemanche, 2020), our data suggests that reduced profilin by *chic* knockdown aggravated defects in cortical actin organization with reduced Dia in *smog* mutants. Overall, our data suggest that Smog is required for cortical actin organization as well as myosin activation during epithelial morphogenesis.

## DISCUSSION

### Smog regulates different pools of myosin in a Fog-dependent and -independent manner during SG invagination

Fog signaling triggers epithelial cell shape changes driving tissue folding and invagination during development of *Drosophila* and other insects (Benton et al., 2019; Manning and Rogers, 2014). During *Drosophila* embryogenesis, *fog* is upregulated in multiple tissues undergoing apical constriction and tissue invagination, such as ventral furrow (mesoderm), the posterior midgut (endoderm), and the SG (ectodermal derivative) (Nikolaidou and Barrett, 2004). During *Drosophila* gastrulation, the ubiquitously expressed GPCR Smog and the mesoderm-specific GPCR Mist respond to Fog signal to regulate myosin contractility during mesoderm invagination (Kerridge et al., 2016; Manning et al., 2013). Here, we provide evidence that Smog functions as a SG receptor for Fog to activate myosin during SG invagination. Knockdown of *smog* by RNAi suppresses the gain-of-function effect of Fog in the SG (Fig. 1). Also, Smog-transfected S2 cells contract upon Fog signal (Fig. 2), suggesting that Smog responds to Fog to regulate myosin contractility both in vivo and in vitro. Our data show that Fog overexpression leads to Smog recruitment to the medioapical region of SG cells (Fig. 1). Consistent with our data, Fog overexpression induces oligomerization of Smog in early *Drosophila* embryos (Jha et al., 2018).

Our study also dissects the roles of Smog in myosin activation during SG morphogenesis and reveals Fog-independent roles of Smog. Whereas *fog* mutants exhibit a decrease in only the medioapical myosin pool in SG cells (Chung et al., 2017; Fig. S3), *smog* loss and SG-specific knockdown of *smog* result in a significant reduction of both medioapical and junctional myosin (Fig. 4). We propose that Smog regulates junctional myosin during epithelial morphogenesis in response to an unknown, ubiquitously expressed ligand and regulates medioapical myosin in response to Fog in tissues with high Fog signals, such as the mesoderm (Kerridge et al., 2016; Manning et al., 2013) and the SG (this study). Consistent with this model, during *Drosophila* gastrulation, Smog activates different myosin pools in tissues with different Fog levels. In the mesoderm, where Fog levels are high, Smog and Mist transduce Fog signal to activate myosin in the medioapical region and drive apical constriction. In the ectoderm, where Fog levels are very low, Smog is required for junctional myosin activation to drive cell intercalation. SG cells undergo both apical constriction and cell intercalation, and our data shows that Smog regulates both pools of myosin in the SG in a Fog-dependent and -independent manner. It is possible that in tissues with high Fog signals, an additional tissue-specific GPCR(s) function with Smog in response to Fog to fine-tune the Fog signaling for proper apical constriction, like Mist does in the mesoderm. It will be interesting to determine whether additional SG-specific GPCRs function with Smog during SG invagination.

The downstream components that transduce signals from the Smog GPCR to regulate myosin in distinct regions of SG cells await discovery. Good candidates include two Rho guanine nucleotide exchange factors (RhoGEFs), RhoGEF2 and Dp114RhoGEF, which have been shown to activate Rho1 signaling at the medioapical and junctional domain of epithelial cells, respectively (Garcia De Las Bayonas et al., 2019). This process requires upstream activation by heterotrimeric G proteins Gα_12/13_/Cta and Gβ13F/Gγ1 at the medioapical and junctional domains, respectively (Garcia De Las Bayonas et al., 2019). Thus, distinct ligand-receptor binding with Smog may activate distinct G proteins and/or RhoGEFs in the medioapical and junctional domains.

### Smog regulates cortical actin organization during SG invagination

We also discover that Smog regulates cortical F-actin organization during SG invagination (Fig. 6). *smog* loss significantly reduces levels of F-actin and the formin protein Dia (Fig. 6). Rho1 regulates actin polymerization via Dia (Goode and Eck, 2007), and apical F-actin formation in the *Drosophila* SG requires apical localization and activity of Dia (Massarwa et al., 2009; Rousso et al., 2013). Our data suggest that Smog activates Rho1 signaling to regulate both cortical F-actin (via Dia) and myosin (via Rok) (Fig. 7). *fog* mutants also show slightly reduced F-actin and Dia levels, to a similar level to *smog* null [mild] embryos (Fig. 7), and these embryos do not show membrane blebbing (Fig. 7; Fig. S5). It is possible that Smog regulates cortical actin in both Fog-dependent and independent manners and perhaps subtle changes in Dia and F-actin levels in *fog* mutants are not enough to cause membrane blebbing. Alternatively, Smog’s role in regulating cortical actin is independent of Fog; reduced F-actin levels in *fog* mutants may be a secondary effect due to a defective medioapical myosin pool. During *Drosophila* gastrulation, *fog* mutants show normal GFP-Moesin signals in the ventral furrow, suggesting normal cortical actin organization upon *fog* loss (Kanesaki et al., 2013). Gα_12/13_/Cta is a key component downstream of Fog that recruits RhoGEF2 to the apical membrane to regulate myosin contractility (Barrett et al., 1997; Dawes-Hoang et al., 2005). Interestingly, mutations in Gαi (G-iα65A) and Gβ13F/Gγ1, but not Gα_12/13_/Cta, disrupt cortical actin organization in early *Drosophila* embryos (Fox and Peifer, 2007; Kanesaki et al., 2013). Therefore, Smog may signal through Gαi and/or Gβ13F/Gγ1 to organize cortical actin, independent of Fog and Gα_12/13_/Cta.

**Figure 7.**
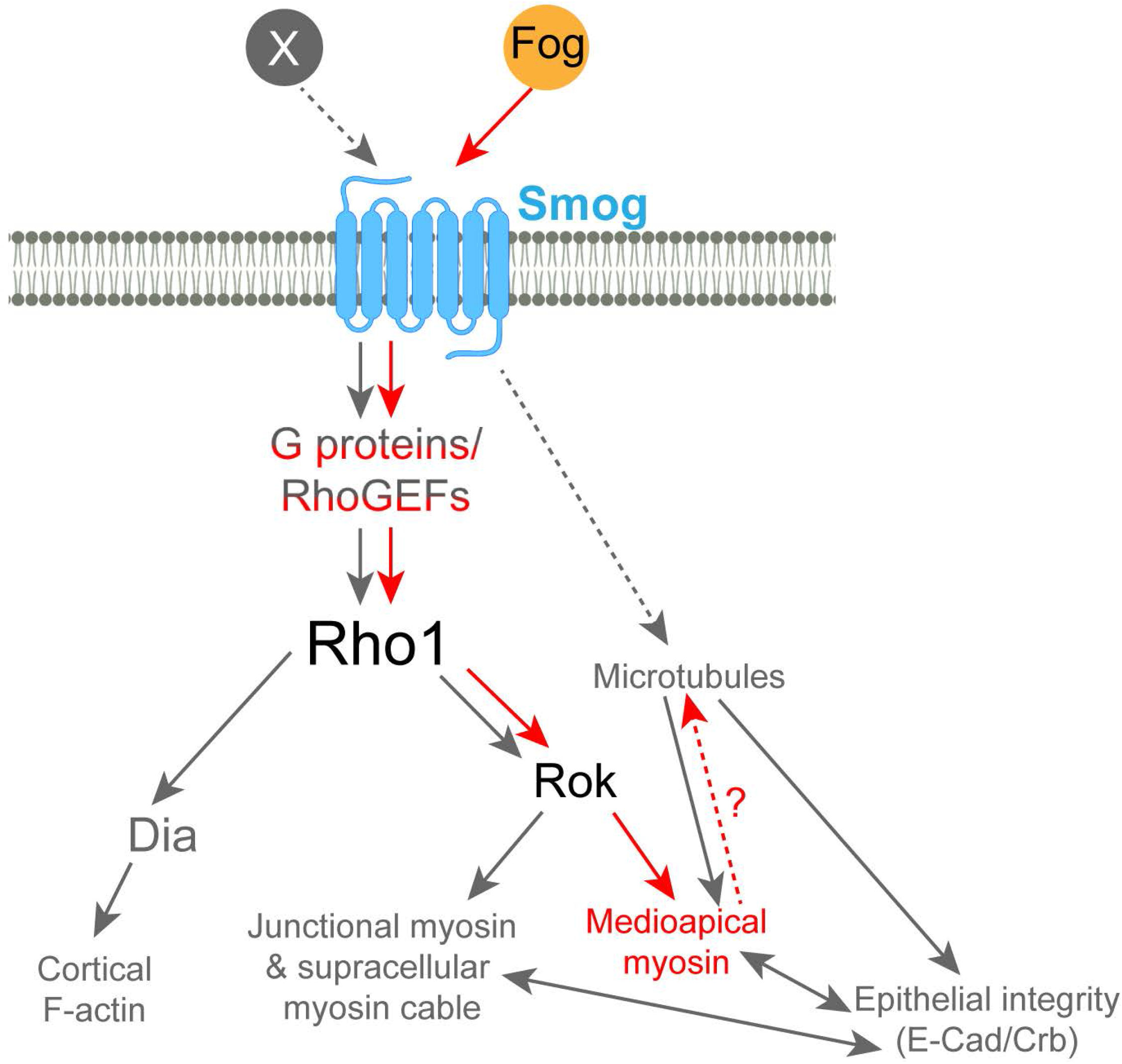
A proposed model for roles of Smog during epithelial tube formation. During SG invagination, Smog responds to Fog signal to promote Rok accumulation and medioapical myosin formation to control apical constriction (red arrows). Independent of Fog (and responding to an as-yet-unknown ligand X), Smog regulates epithelial integrity by regulating junctional and the supracellular myosin pools, the microtubule networks, and key apical/junctional components (gray arrows). Smog is also required for cortical actin organization during SG invagination, through regulating Dia levels (gray arrows). Different ligands may recruit specific G proteins and RhoGEFs to activate distinct downstream effectors.

The link between heteromeric G proteins and Rho1 signaling occurs through activation of specific RhoGEFs (Vázquez-Victorio et al., 2016). Actin polymerization resulting from Fog-independent Smog activation may imply the recruitment of distinct proteins modulating the activation of Rho1-mediated pathways. For example, in the absence of Fog, a specific RhoGEF(s) downstream of Gαi or Gβ13F/Gγ1 may link Smog and Rho1 signaling to regulate cortical actin organization; upon high Fog signals, a different Rho1 modulator(s) downstream of Gα_12/13_/Cta may be recruited to facilitate medioapical myosin formation. Further work is needed to identify the link between Smog and Rho1 activation in the presence or absence of Fog during SG invagination. It will be also interesting to determine whether Fog-independent roles of Smog–regulating cortical actin and junctional myosin–involve common or distinct Rho1 modulators.

### Smog regulates epithelial integrity during development

*smog* null mutant embryos show a range of defects, allowing us to dissect the roles of Smog in epithelial morphogenesis. Our data reveal that Smog is required for epithelial integrity by affecting levels and localization of Crb and E-Cad (Fig. 5). Since the decrease in the level of Crb and E-Cad in *smog* mutants correlates with significantly reduced myosin levels, disrupted epithelial integrity may be due to disrupted actomyosin networks. Myosin II is critical for cells to concentrate E-cad at cell-cell contacts (Shewan et al., 2005). In early *Drosophila* embryos, the absence of contractile myosin leads to the disassembly of junctions (Weng and Wieschaus, 2016). A possible scenario is that Smog is required to recruit key Rho1 regulators, such as RhoGEF2, Dp114RhoGEF, or other SG-upregulated RhoGEFs, to AJs during SG invagination to promote junctional myosin assembly. Since Dia helps coordinate adhesion and contractility of actomyosin (Homem and Peifer, 2008) and formin-mediated actin polymerization at AJs stabilizes E-Cad and maintains epithelial integrity (Rao and Zaidel-Bar, 2016), reduced Dia levels in *smog* mutants may also contribute to loss of epithelial integrity.

Reduced and discontinuous signals of apical and junctional components in *smog* mutant embryos can also be due, in part, to compromised intracellular trafficking. Disorganized microtubule networks in SGs in *smog* mutants (Fig. 5) support this idea. Our recent study showed that microtubule- and Rab11-dependent intracellular trafficking regulates apical myosin pools and apical constriction during SG invagination, via apical enrichment of Fog and the continuous distribution of Crb and E-Cad along junctions (Le and Chung, 2021). The mechanism of how Smog affects microtubule organization remains to be revealed. *fog* mutants also show slightly disrupted microtubule networks, albeit less prominently than *smog* null [severe] (Fig. 5). Whether this is a direct effect of loss of *fog* or an indirect result of the disrupted medioapical pool of myosin remains to be answered. Overall, our findings suggest the multifaceted roles of Smog during epithelial tube formation in regulating distinct myosin pools and cortical actin in different subcellular domains in a ligand-dependent manner.

## MATERIALS AND METHODS

### Fly stocks and genetics

The fly stocks used in this study are listed in Table S1. All the crosses were performed at 25°C.

### Immunofluorescence and confocal microscopy

*Drosophila* embryos were collected on grape juice agar plates supplemented with yeast paste at 25 °C. Embryos were dechorionated with 50% bleach. For most samples, embryos were fixed in 1:1 heptane: formaldehyde for 40 minutes and devitellinized with 80% ethanol. For Rok-GFP, sqh-GFP, and phalloidin staining, embryos were hand-devitellinized. Embryos were then stained with primary and secondary antibodies in PBSTB (1X PBS with 0.1% Triton X-100 and 0.2% bovine serum albumin) and PBTr (1X PBS with 0.1% Triton X-100) for ethanol devitalization and hand-devitalization, respectively. Antibodies used in our experiments are listed in Table S2. Embryos were mounted in Aqua-Poly/Mount (Polysciences, Inc) and imaged with a Leica TCS SP8 confocal microscope using 63x, NA 1.4 and 40x, NA 1.3 objectives. Images were acquired as z-stacks (each 0.3 μm apart) that span the apical and junctional domains of cells in the SG placode.

### Fluorescent in situ hybridization

Embryos were fixed using formaldehyde/heptane for 30 min followed by devitellinization with methanol. SP6 polymerase-synthesized digoxigenin (DIG)-labeled antisense probe was prepared using the following primers: *smog*-F, 5’-ACAGAGCCCACCTGTGTAGG- 3’; *smog*-R, 5’-TCGCTGATCGAAAATGATCTC-3’. Fluorescent in situ hybridization was performed using standard methods as described in (Knirr et al., 1999). Briefly, the embryos were pre-hybridized for 1 hour at 56°C post-fixation. Following this step, the embryos were hybridized with the probe overnight at 56°C. The next day, embryos were stained for DIG using the anti-DIG primary antibody and biotin-conjugated secondary antibody. The embryos were then incubated with AB solution (PK-6100, VECTASTAIN Elite ABC kit). Tyramide signal amplification reaction was performed using Cy3 fluorescent dye (diluted 1:50 using amplification diluent) (Perkin Elmer Life Sciences, Inc., NEL753001kt). The embryos were co-stained with CrebA and Crb antibodies for visualizing SG nuclei and apical cell boundaries, respectively.

### Total Smog-GFP and Dia intensity

For Smog-GFP quantification, maximum intensity projections of two z-sections that span the apical and the junctional region of SG cells were used. Intensity means of Smog-GFP signals of SG cells in the entire SG placode were measured using Imaris software (Bitplane). The mean intensity of Smog-GFP was normalized by the median deviation. The integrated density of Smog-GFP was calculated by multiplying the apical area of the cell with the mean intensity of each cell. Total Smog-GFP signals within the whole placode were calculated as the sum of the integrated density of all cells. For background correction, mean gray values of Smog-GFP in ten cells outside of the SG placode were measured using ImageJ (NIH). The average value of mean gray values of Smog-GFP in these ten cells was used to subtract the background of the cells inside the placode from the same embryo. Five SGs were used for quantification for each genotype except for control in Fig. 1E, where four SGs were used. Statistical significance was determined using Mann-Whitney U test in the GraphPad Prism software.

For total Dia intensity quantification, maximum intensity projection of two z-sections of the most apical region of SG cells was used. The images were then exported to ImageJ and converted to the RGB stack image type. Mean gray values of Dia signals in the whole placode were measured. For background subtraction, the average mean gray value of Dia signals for ten cells outside the SG placode was subtracted from the mean gray value of Dia intensity of the whole SG placode. These values were scaled by the mean normalization. The total Dia intensity in the whole placode was calculated by multiplying the normalized intensity with the entire area of the placode. Statistical significance was determined using Mann-Whitney U test.

### Cell segmentation and apical area quantification

SGs that were invaginated within the range of 4.8-9.9 μm depth were used for quantification. Two or three z-sections of the apical domain were used to generate a maximum intensity projection (Leica LasX software). SG cells were marked using CrebA and segmented along E-Cad signals using the Imaris program. Apical areas of segmented cells were calculated using Imaris, and cells were color-coded based on their apical domain size. Frequency distribution was performed using GraphPad Prism. Apical areas from eight SGs were quantified in control, *smog* knockdown (Z), and *smog* null [mild] embryos. Six SGs were used for quantification for *smog* knockdown (M/Z) and *smog* null [severe] embryos. Statistical significance was determined using Mann-Whitney U test (percentage of cells) and Kolmogorov-Smirnov test (cumulative percentage of cells).

### Quantification of myosin intensity in the medioapical and junctional domains

Maximum intensity projection of two z-sections spanning apical and junctional regions of SG cells with sqh-GFP and E-Cad signals was used. The images were then exported to ImageJ and converted to RGB stack image type. Twenty cells near the invagination pit in each of five SGs were used for quantification. To calculate the medioapical and junctional myosin intensity, regions were drawn manually along the inner and outer boundary of the E-Cad signal of each cell, and mean gray values were measured for medioapical and junctional myosin. For background subtraction, the average mean gray value of sqh-GFP signals for ten cells outside the SG placode was subtracted from the mean gray value of medioapical and junctional myosin for each SG cell. SuperPlots (Lord et al., 2020) were used to address the variability of datasets in each SG. Each data point in the graph represents one SG consisting of twenty cells used for quantification. Statistical significance was determined using Welch’s t-test. A normal distribution was tested using Jarque-Bera test (Microsoft Excel) or a bell-shape of data distribution by making a histogram.

### Quantification of areas of Rok-GFP and medioapical myosin particles

A single z-section of the confocal image with the strongest Rok-GFP signals in the medioapical region was used. Rok-GFP signals were converted to black and white using the threshold tool in Adobe Photoshop. Junctional Rok-GFP signals were removed manually based on E-Cad signals. Areas of Rok-GFP puncta were determined using the Analyze Particles tool in ImageJ. Rok-GFP puncta with area ≥ 0.02 μm^2^ was measured. Fifteen cells in the dorsal posterior region near the invagination pit of the SG placode were used for quantification. The same strategy was used for sqh-GFP signals to quantify the area of medioapical myosin. SuperPlots show mean values of data points of five SGs. Statistical significance was determined using Mann-Whitney U test.

### Quantification of the waviness of junctions

Using a single z-section with the highest E-Cad signals and ImageJ software, the shortest (L_0_) and the actual distance (L) between vertices were measured in SG cells. The ratio of these distances (L/L_0_) was used as the waviness of junctions. 10-15 cells near the invagination pit in each of five SGs were used for quantification. Statistical significance was determined using Mann-Whitney U test.

### Circularity of the SG boundary

Using confocal images and ImageJ, cell boundaries were manually drawn along E-Cad signals at the SG placode boundary and one cell row outside and inside the placode. For most samples, strong myosin cable signals at the dorsal, anterior, and posterior boundaries of the placode and CrebA signals in SG cells were used to determine the SG boundary. In *smog* null [severe], where myosin signals were significantly reduced in all cells, CrebA signals were used to determine the boundary. The ventral midline was used as the ventral boundary of the SG in all cases. The perimeter and the area of the SG corresponding to these boundaries were measured using ImageJ, and circularity was calculated using the formula, C = 4π area/perimeter^2^. For a perfect circle, the value of circularity should be 1. Five SGs were used for quantification, and statistical significance was determined using Welch’s t-test.

### Quantification of number and length of gaps of Crb and E-Cad

The number and the length of gaps for Crb and E-Cad signals were measured using confocal images and ImageJ. A single z-section with the highest Crb or E-Cad signals was chosen for quantification. Gaps with a length ≥0.2 μm were used for quantification. All junctions from ten cells in five SGs were quantified. If there were multiple gaps on a\ junction, we added the length of all the gaps on that junction to calculate the ratio of the length of gaps to junctional length. Statistical significance was determined using Mann-Whitney U test.

### Acetylated and tyrosinated α-tubulin intensity quantification

Three z-sections spanning the most apical region of SG cells were used for quantification. The SG boundary was drawn manually based on the E-Cad and CrebA signals. The mean gray value of Ace-tub signals in the whole SG placode and ten cells outside the SG of the same embryo was measured using ImageJ. The ratio of the mean gray value of Tyr-tub signals inside the placode to the average mean gray value of cells outside the placode was calculated and plotted. The same method was used for Tyr-tub. Five SGs for each genotype were used for quantification. Mann-Whitney U test was used to calculate the p values.

### Cell contractility assay

cDNA clones for *smog* (RE70685) and *mist* (RE13854) were obtained from *Drosophila* Genomics Resource Center (DGRC). Isolated cDNAs were cloned in-frame with the V5 tag into the pMT-V5-HisB vector (Addgene) using the conventional restriction digestion and ligation method. Primers with restriction sites used to isolate the open reading frame are stated below (restriction sites underlined).

*smog* CDS-EcoRI-5’, 5’-CCG<u>GAATTC</u>ATGGAACTGTGCATAGCAAC-3’

*smog* CDS-NotI-3’, 5’-ATTT<u>GCGGCCGC</u>ATTGGTCGTGATTGTATCTTTGG-3’

*mist* CDS-EcoRI-5’, 5’-CCG<u>GAATTC</u>ATGGACAGGAGTCGGAGTAGC-3’

*mist* CDS-NotI-3’, 5’-ATTT<u>GCGGCCGC</u>AGCAAATGGTCTCCATTTTG-3’

S2 and S2-Mt-Fog-myc cells (Manning et al., 2013) were obtained from DGRC. Cells were tested for contamination regularly. Shields and Sang M3 insect media (HiMedia) was used with 10% Fetal Bovine Serum (Corning) and 1:1000 Antibiotic-Antimycotic (Gibco). Cells were cultivated at 25°C either in 25-cm^2^ flask in 5 ml media or 75-cm^2^ flask in 15 ml media. To prepare for the Fog media, S2-Mt-Fog-myc cells were grown undisturbed for 4-8 days to attain approximately 100% confluency. 50 μl of 100 mM CuSO4 was added to nearly 100% confluent cells in 75-cm^2^ flask to induce metallothionein promoter. The Fog-containing media was collected by centrifugation and further concentrated using 3,000 MW concentrators. The presence of Fog was confirmed by Western blot using the Myc-antibody (Invitrogen; RRID:AB_2533008). Non-transfected S2 cells were used as a control.

Cell contractility assay was performed as described in (Manning et al., 2013). Smog- or Mist-transfected S2 cells were induced with CuSO_4_. Cells were then transferred to Concanavalin A-coated coverslips and fixed using 10% paraformaldehyde in PBTr for 15 minutes at room temperature. After being blocked in 5% NGS in PBTr for 20 minutes at RT, cells were stained with the V5 antibody (Invitrogen; RRID:AB_2556564) to differentiate between transfected and non-transfected cells. Cell contractility of transfected cells was monitored using a phase-contrast filter in a Leica DM2500 microscope using the 40X, 0.8 NA objective. Three independent experiments were performed. Two-way ANOVA was performed to calculate statistical significance.

## Acknowledgments

We thank the members of the Chung laboratory for their comments and suggestions. We thank A. Martin, T. Lecuit and the Bloomington stock center for fly stocks, and D. J. Andrew and the Developmental Studies Hybridoma Bank for antibodies. We thank Flybase for the gene information. We are grateful to D. J. Andrew, C. D. Hanlon, J. Kim, and J. Matthew for their helpful comments on the manuscript. This work is supported by a start-up fund from Louisiana State University and grants from the Louisiana Board of Regents Research Competitiveness Subprogram and the National Science Foundation (MCB 2141387) to S.C.

