## Supplemental figures, legends, tables for "Multifunctional role of GPCR signaling in epithelial tube formation"

### Sup Fig. S1

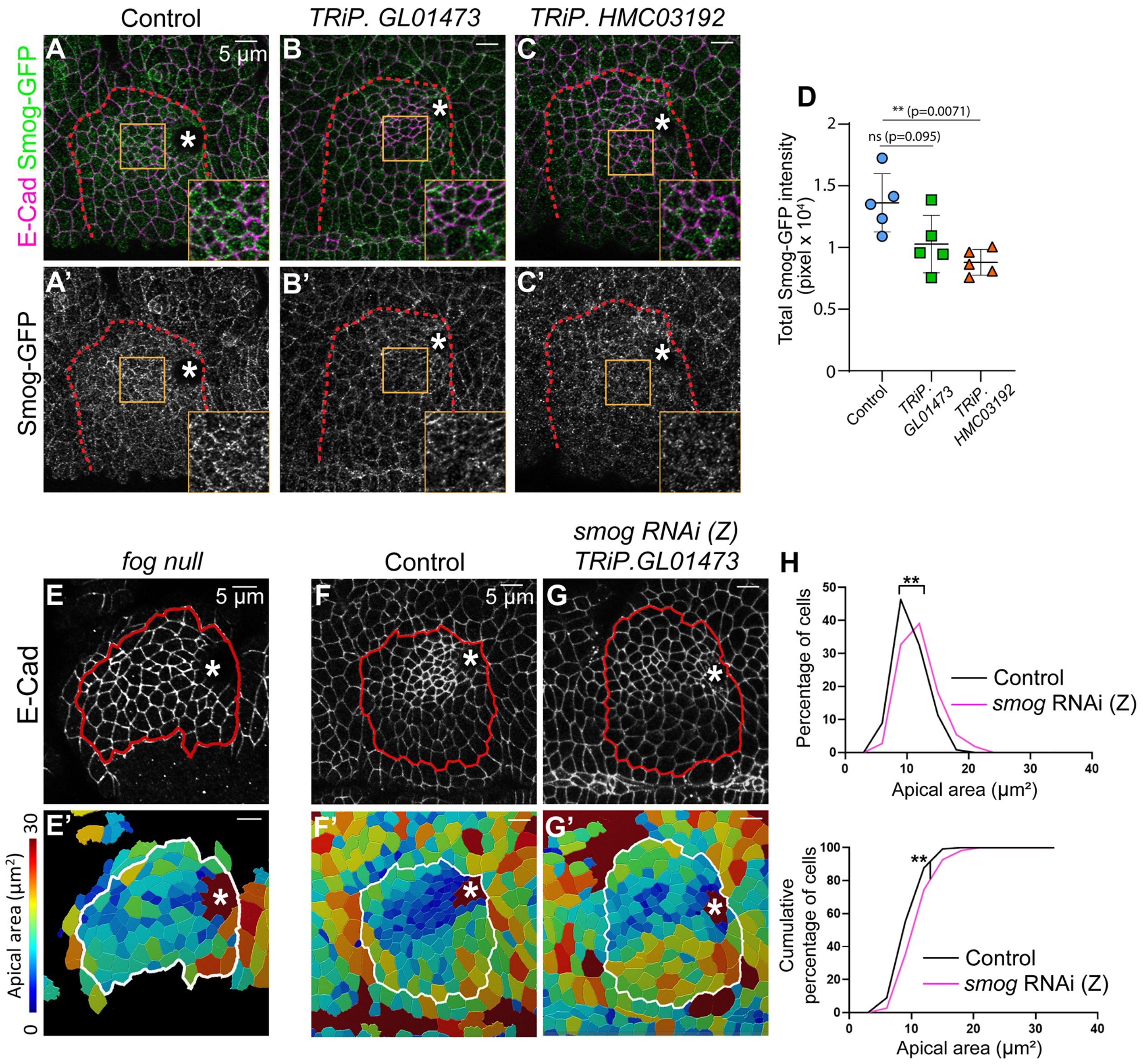


**Supplementary Figure S1. Verification of *smog* knockdown using two RNAi lines and apical constriction defects with a weaker line.** (A-C’) E-Cad (magenta) and Smog-GFP (green) signals in stage 11 SGs in control (A, A’), *smog* zygotic knockdown using *TRiP.GL01473* (B, B’) and *TRiP.HMC03192* (C, C’). Insets, magnified view of yellow boxed regions in the SG. (D) Quantification of total Smog-GFP intensity in SGs. n=5 SGs for each genotype. ns, non-significant; **p≤0.01 (Mann-Whitney U test). (E, E’) Confocal image (E) of the SG in *fog* mutants and the corresponding heat map of apical areas (E’). The embryo surface is often uneven in *fog* mutant embryos due to additional folding and grooves, resulting in some cells outside the SG being out of focus and not shown in the image. (F-G’) Confocal images of SGs in control (F) and zygotic knockdown of *smog* by using TRiP.GL01473 (G) and corresponding heat maps of apical areas (F’ and G’). Red and white lines, SG boundary. Asterisks, invagination pit. (H) Quantification of percentage and cumulative percentage of cells with different apical areas. Mann-Whitney U test (for percentage of cells) and Kolmogorov-Smirnov test (for cumulative percentage of cells). n=8 SGs (control, 902 cells; smog RNAi (Z), 856 cells). $^{**}p \leq 0.01$.

**Sup Fig. S2**

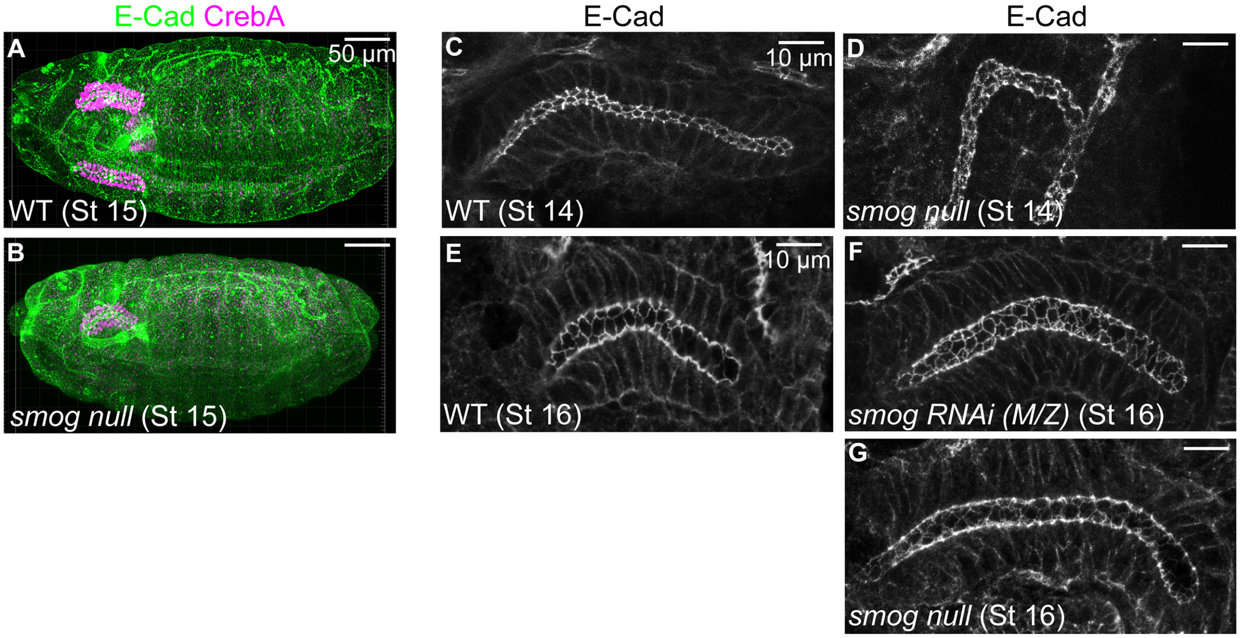


**Supplementary Figure S2. SGs in *smog* null mutants have relatively normal morphology at stage 14 and later during embryogenesis.** (A, B) Confocal images of stage 15 WT (A) and *smog* null embryos stained for E-Cad (green) and CrebA (magenta). (C-G) SGs stained for E-Cad in WT (C, E), *smog* M/Z knockdown (F), and *smog* null mutant (D, G) embryos. All embryos at stage 14 and later have relatively normal morphology, including the SG, except for a rare case of the crooked SG in *smog* null mutants (D).

Supp Fig. 3

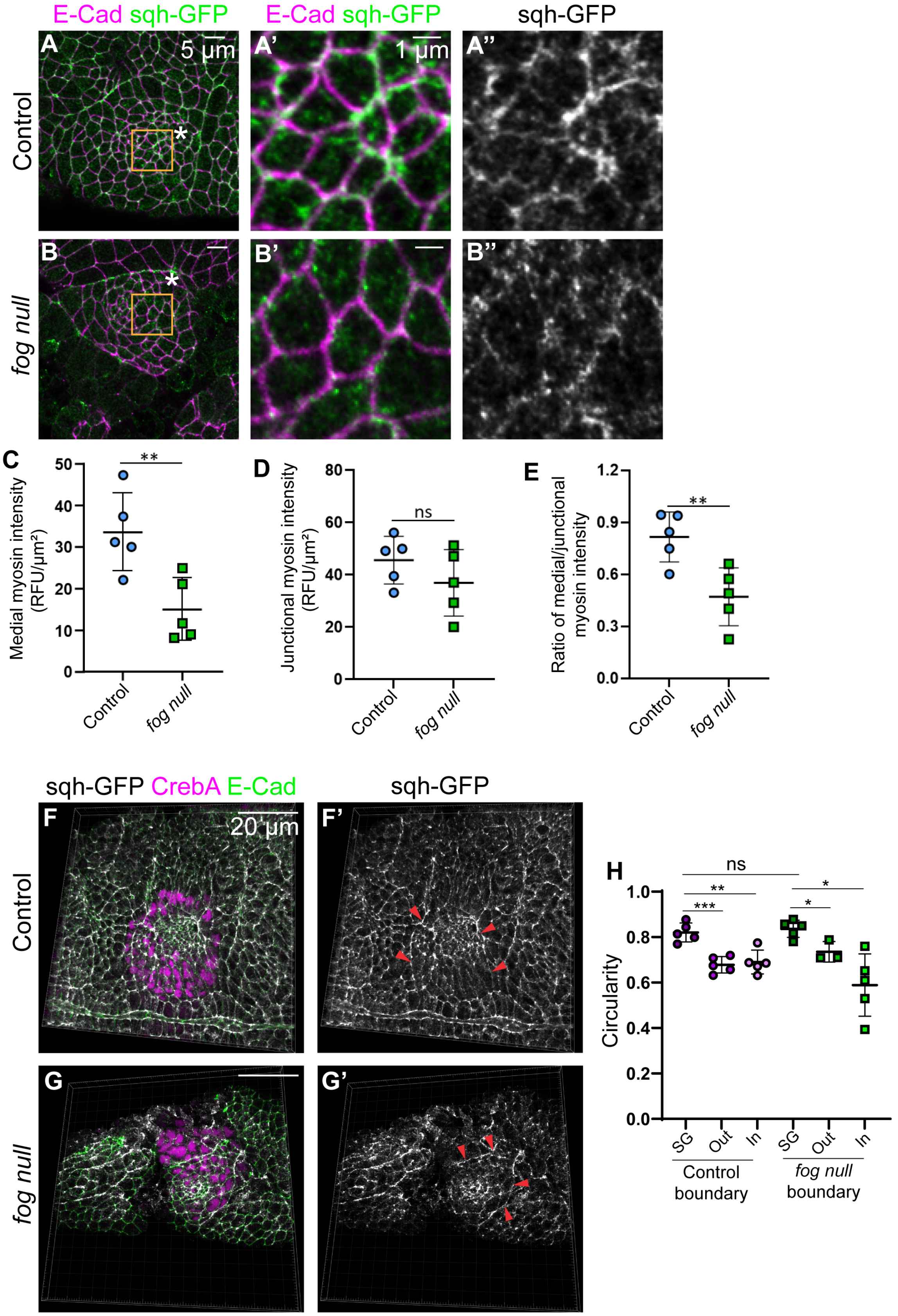

E-Cad sqh-GFP
E-Cad sqh-GFP
sqh-GFP
A
5 µm
A'
1 µm
A''
Control
B
B'
B''
fog null
C
Medial myosin intensity (RFU/µm²)
**
Control
fog null
D
Junctional myosin intensity (RFU/µm²)
ns
Control
fog null
E
Ratio of medial/junctional myosin intensity
**
Control
fog null
sqh-GFP CrebA E-Cad
sqh-GFP
F
20 µm
F'
Control
G
G'
fog null
H
Circularity
ns
**
***
*
*
SG
Out
In
SG
Out
In
Control boundary
fog null boundary

**Supplementary Figure S3. Fog is required for proper medioapical myosin formation.** (A-B’’) Confocal images of stage 11 SGs with sqh-GFP (green) and E-Cad (magenta) signals. Compared to control (A-A’’), sqh-GFP signals are reduced in medioapical regions SGs with *fog* null embryos (B-B’’). (A’-B’’) Higher magnification of the yellow boxed region in A-B. (C-E) Quantification of medioapical myosin intensity (C), junctional myosin intensity (D), ratio of medial and junctional myosin intensity (E). n=5 SGs, 100 cells for each genotype. **$p \leq 0.01$ (Welch’s t-test). (F, G) 3D reconstruction of stage 11 SGs stained with E-Cad (green), sqh-GFP (white), and CrebA (magenta) in control (F) and *fog* null mutant embryos (G). The supracellular myosin cable surrounds the entire SG placode in WT and *fog* mutant embryos (red arrowheads in F’ and G’). (H) Quantification of the circularity as a measure of smoothness and tension of the SG, outer, and inner boundary in control and *fog* null SGs. n=5 SGs for each genotype. *$p \leq 0.05$; **$p \leq 0.01$; ***$p \leq 0.001$ (Welch’s t-test).

### Sup Fig. S4

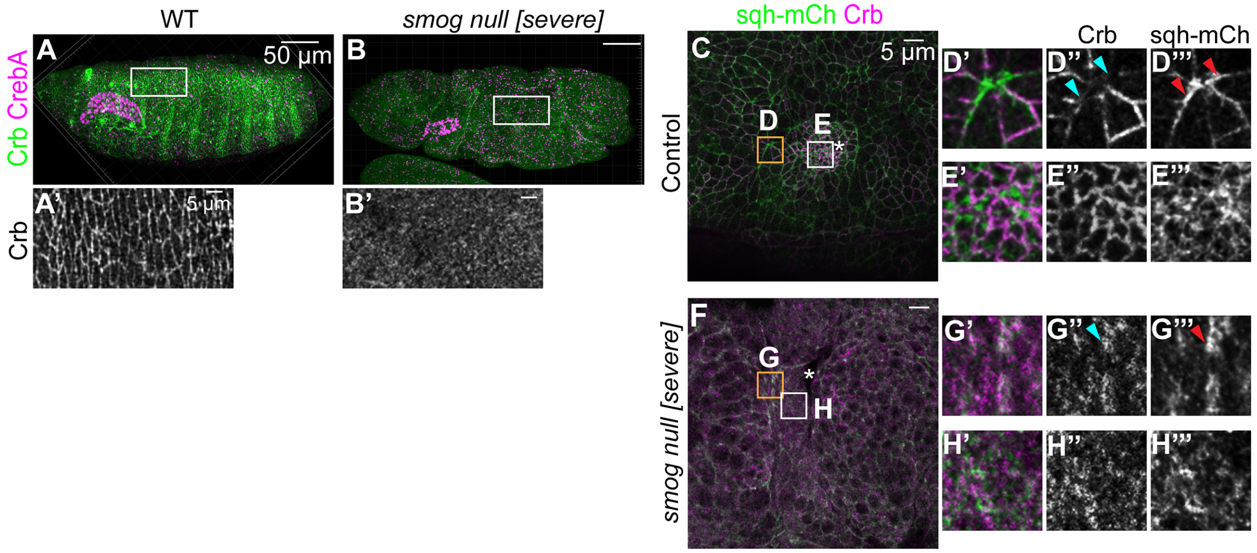


**Supplementary Figure S4. *smog* loss leads to the reduction of Crb levels in the whole embryo and loss of Crb anisotropy at the SG boundary.** (A, B) Confocal images of stage 14 WT (A) and *smog* null [severe] mutant (B) embryos stained for Crb (green) and CrebA (magenta). (A', B') Magnified view of the white boxed region in A and B. Compared to clear Crb signals at the cell boundary in WT (A'), Crb levels are reduced and diffused in some *smog* null mutant embryos with a severely defective morphology (B'). (C-H''') control (C-E''') and *smog* null [severe] embryos (F-H''') showing sqh-mCh (green) and Crb (magenta) signals. (D, G) Cells at the SG boundary. (E, H) SG cells near the invagination pit (asterisk). (D'-H''') Higher magnification of the boxed regions. In contrast to high myosin (red arrowheads in D''') and low Crb (cyan arrowheads in D'') levels in the SG boundary in control embryos, the overall Crb levels are reduced in *smog* null embryos, showing loss of anisotropic localization of Crb at the SG boundary (G-H''').

### Sup Fig. S5

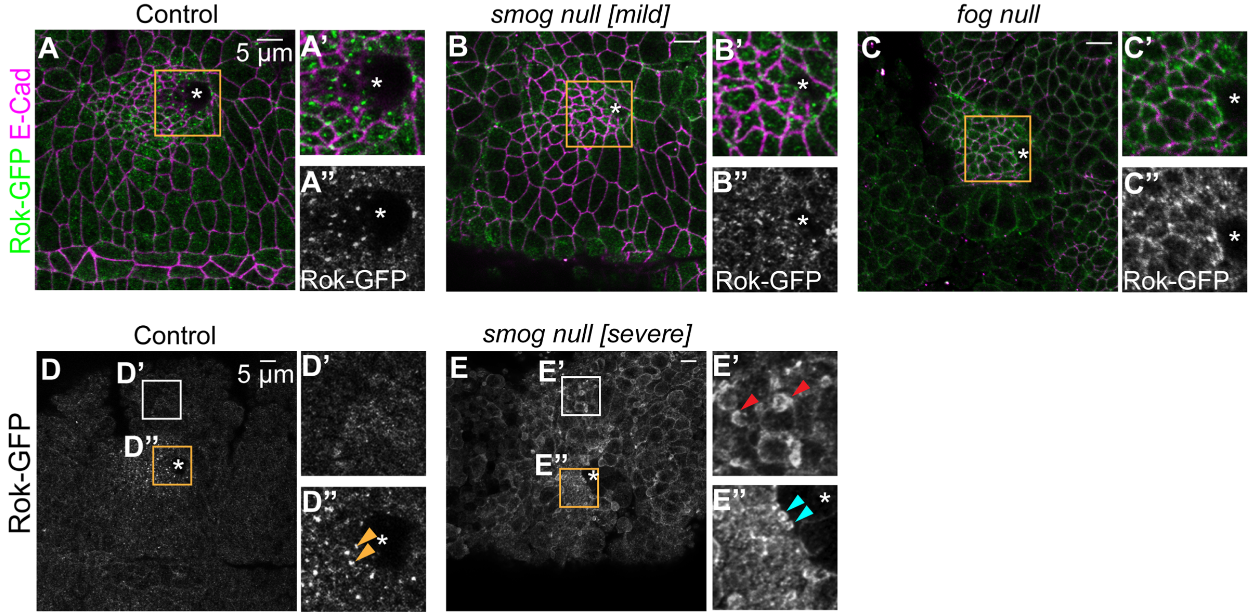


**Supplementary Figure S5. *smog* loss leads to bleb formation in SG cells as well as outside the SG placode.** (A-C'') Confocal images for Rok-GFP (green) and E-Cad (magenta) signals in stage 11 SGs in control (A-A''), *smog* null [mild] (B-B''), and *fog* mutant (C-C'') embryos. Blebs are not observed in these embryos. Due to the uneven embryo surface in *fog* mutants, apical/junctional and more basal regions are shown for SG cells near and far from the invagination pit, respectively, in the single z-section. (D-E'') Confocal images for Rok-GFP signals in control (D-D'') and *smog* null [severe] (E-E'') embryos. Blebs are observed both in non-SG cells outside the SG placode (red arrowheads in E') and SG cells (cyan arrowheads in E''). Yellow arrowheads in D'', normal Rok accumulation in the medioapical region of SG cells. Asterisk, invagination pit.

**Table S1. Fly strains used**

| Fly strain | Sources and References | RRID |
|---|---|---|
| *Oregon R* (wild type) | | |
| *fkh-Gal4* | Henderson and Andrew, 2000 | BDSC_78060 |
| *matα4-Gal4* (on II) | Bloomington Stock Center | BDSC_7062 |
| *matα4-Gal4* (on III) | Bloomington Stock Center | BDSC_7063 |
| *UAS-Fog* | Dawes-Hoang et al., 2005 | |
| *smog-GFP* | Kerridge et al., 2016 | |
| *ubi-Rok-GFP* | Abreu-Blanco et al., 2014 | BDSC_52289 |
| *sqh-GFP* | Royou et al., 2004 | BDSC_57144 |
| *sqh-mCherry* | Martin et al., 2009 | BDSC_59024 |
| *smog* null mutant | Kerridge et al., 2016 | |
| $fog^{RA67}$ | | BDSC_6218 |
| $fog^{S4}$ | | DGGR_106665 |
| *UAS-Dicer-2* | Vienna *Drosophila* Resource (#60008) | |
| *TRiP.GL01473* (*smog* RNAi (weak)) | Bloomington Stock Center | BDSC_43135 |
| *TRiP.HMC03192* (*smog* RNAi (strong)) | Bloomington Stock Center | BDSC_51705 |
| *TRiP.HMS00550* (*chic* RNAi) | Bloomington Stock Center | BDSC_34523 |

**Table S2. Antibodies Used**

| Antibody | Source | RRID | Dilution |
|---|---|---|---|
| α-E-Cad (rat) | DSHB (DCAD2) | AB_528120 | 1:50 |
| α-CrebA (rat) | Andrew lab (John Hopkins University); This study | | 1:3000 |
| α-CrebA (rabbit) | Andrew lab (John Hopkins University) | AB_10805295 | 1:5000 |
| α-CrebA (guinea pig) | This study | | 1:3000 |
| α-GFP (chicken) | Invitrogen (A10262) | AB_2534023 | 1:500 |
| α-β-galactosidase (rabbit) | Invitrogen (A11132) | AB_221539 | 1:500 |
| α-β-galactosidase (mouse) | Invitrogen (MA5-15222) | AB_10980770 | 1:500 |
| α-Crb (mouse) | DSHB (Cq4) | AB_528181 | 1:10 |
| α-mCherry (rat) | Invitrogen (M11217) | AB_2536611 | 1:1000 |
| α-mCherry (rabbit) | Invotrogen (PA534974) | AB_2552323 | 1:1000 |
| α-Myc (mouse) | Invitrogen (13-2500) | AB_2533008 | 1:250 |
| α-V5 (mouse) | Invitrogen (R960-25) | AB_2556564 | 1:400 |
| α-acetylated α-tubulin (mouse) | Invitrogen (32-2700) | AB_2533073 | 1:1000 |
| α-tyrosianted α-tubulin (rat) | Invitrogen (MA1-80017) | AB_2210201 | 1:1000 |
| α-Dia (rabbit) | Afshar et al., 2000 | | 1: 5000 |

| | | | |
|---|---|---|---|
| Phalloidin Alexa 488 | Invitrogen (A12379) | | 1:250 |
| Alexa Fluor 488/568/647- coupled secondary antibodies | Invitrogen | | 1:500 |
